# Defined factors to reactivate cell cycle activity in adult mouse cardiomyocytes

**DOI:** 10.1101/755991

**Authors:** Justin Judd, Jonathan Lovas, Guo N. Huang

## Abstract

Adult mammalian cardiomyocytes exit the cell cycle during the neonatal period, commensurate with the loss of regenerative capacity in adult mammalian hearts. We established conditions for long-term culture of adult mouse cardiomyocytes that are genetically labeled with fluorescence. This technique permits reliable analyses of proliferation of pre-existing cardiomyocytes without complications from cardiomyocyte marker expression loss due to dedifferentiation or significant contribution from cardiac progenitor cell expansion and differentiation in culture. Using this system, we took a candidate gene approach to screen for fetal-specific proliferative gene programs that can induce proliferation of adult mouse cardiomyocytes. Using pooled gene delivery and subtractive gene elimination, we identified a novel functional interaction between E2f Transcription Factor 2 (E2f2) and Brain Expressed X-Linked (Bex)/Transcription elongation factor A-like (Tceal) superfamily members Bex1 and Tceal8. Specifically, Bex1 and Tceal8 both preserved cell viability during E2f2-induced cell cycle re-entry. Although Tceal8 inhibited E2f2-induced S-phase re-entry, Bex1 facilitated DNA synthesis while inhibiting cell death. In sum, our study provides a valuable method for adult cardiomyocyte proliferation research and suggests that Bex family proteins may function in modulating cell proliferation and death decisions during cardiomyocyte development and maturation.

## INTRODUCTION

Human and mouse cardiomyocytes (CMs) undergo endoreduplication shortly after birth followed by cell cycle exit, resulting in mostly polyploid mature cardiomyocytes (1–5). Neither species exhibits significant proliferation of cardiomyocytes in adulthood, potentially explaining why adult mammalian hearts do not regenerate. Neonatal mice (before postnatal day 7) and lower vertebrates (through adulthood) have demonstrated robust heart regeneration, due to the dedifferentiation and proliferation of pre-existing differentiated cardiomyocytes (6,7). However, as mouse cardiomyocytes undergo maturation in the neonatal period, they lose the ability to proliferate and so does the adult mammalian heart lose the ability to regenerate (6). Indeed, the rarity of myocardial-derived cancers (8,9) is consonant with the lack of proliferative potential of adult mammalian cardiomyocytes. Thus, the mechanism of permanent cell cycle exit and the potential for induced proliferation of mature mammalian cardiomyocytes has been a topic of intense investigation (10–13) with potential for therapeutic intervention in cardiac diseases involving loss of cardiomyocytes.

Several articles in the last decade have reported increased cell cycle activity through the overexpression or suppression of various purported regulators of the cardiomyocyte cell cycle, reviewed elsewhere (14–18). Suppression of the Hippo pathway with constitutively active Yap1 increased cardiomyocyte proliferation without increasing cardiomyocyte size (10), and improved heart function after myocardial infarction (19). Later work showed Yap interacts with Pitx2 to upregulate antioxidants, which improved recovery after heart injury (20). Still, the complex role of Hippo signaling is unraveling, and has been implicated in cardiomyocyte survival, as well as cardiac tissue remodeling and hypertrophic adaptation in heart failure (21–23). Tbx20 overexpression in vivo increased cardiomyocyte proliferation and reduced expression of senescence marker p16Ink4a, one of two proteins (the other being p19ARF/p14ARF in mice/humans, respectively) encoded by the Cdkn2A locus. Recently, Tbx6 was identified as a single factor that could increase cell cycle activity in postnatal and adult rat cardiomyocytes (24). Silencing of a long non-coding RNA, cardiomyocyte proliferation regulator (CPR) (25), or suppression of miRNA 128 (26) was found to increase cardiomyocyte cell cycle activity and help restore function after myocardial injury. Amazingly, four factors (Cdk1, Cdk4, Cyclin B1, Cyclin D1) were sufficient to drive post-mitotic cardiomyocytes through cytokinesis and improve myocardial function post-infarction (27). Downregulation of Meis1 was shown to increase cardiomyocyte proliferation and was later found to play a role in the switch from glycolytic to oxidative metabolism (28), a key event in the maturation of cardiomyocytes driven in large part by thyroid signaling (29).

Collectively, there seem to be many potential proteins that can stimulate re-entry of CMs into the cell cycle. In this work, we used an in vitro screen to identify novel factors that can contribute to CM proliferation. However, the study of cardiomyocyte proliferation in vitro using fixed cell imaging is limited when the cells of interest dedifferentiate and lose marker identification. Wang et al. developed an adult mouse cardiomyocyte culture using lineage traced cardiomyocytes to demonstrate that adult mouse CMs require dedifferentiation in order to regain a proliferative phenotype (Wang et al. 2017). CMs were shown to lose cardiac troponin I during dedifferentiation in vitro, but could be identified using the genetic fate mapped marker Myh6-MerCreMer-dTomato.

Here, we established a long-term adult mouse cardiomyocyte culture to enable genetic screening using well-characterized lineage tracing mouse genetics for facile identification of dedifferentiating cardiomyocytes. A candidate gene library was constructed by comparing novel microarray data from proliferative embryonic hearts and non-proliferative adult hearts. The candidate gene pool was further enriched for driver genes by selecting against genes downregulated in adult epicardium, and then supplemented with genes mined from published RNASeq and microarray data, creating the final candidate gene library. Each candidate gene was packaged into an adenovirus vector and delivered to cardiomyocytes en masse.

Cell cycle analysis by 5-ethynyl-2’-deoxyuridine (EdU) labeling revealed a high percentage of S-phase re-entry by overexpression of the entire pool of candidate genes. Subtractive pools identified known cell cycle protein E2f2 as sufficient for S-phase re-entry in cultured adult mouse cardiomyocytes. However, in contrast to previous work in neonatal mouse cardiomyocytes (30,31), we found that overexpression of E2f2 caused massive cell death in adult mouse cardiomyocytes when cultured for several days after gene delivery. TUNEL staining showed elevated DNA nicking in E2f2-treated cultures, suggesting apoptosis may be partially responsible for cell death. Since the full pool of candidate genes did not cause widespread death, we then performed an additional screen to identify factors in the library that rescue cell viability. From this screen, we found two proteins from the Brain Expressed X-Linked (Bex)/Transcription elongation factor A-like (Tceal) superfamily, Tceal8 and BEX1, that rescue cell viability during E2f2 ectopic expression. Although both proteins rescued cell viability, Tceal8 inhibited S-phase re-entry in the presence of E2f2, while BEX1 allowed E2f2-induced S-phase re-entry. Therefore, this study implicates Bex1 in the control of cell death decisions during cardiomyocyte replication and maturation.

## RESULTS

### Adult Mouse Cardiomyocyte Isolation and Culture

We developed a long-term adult mouse cardiomyocyte culture using well-characterized lineage tracing mouse genetics for facile identification of dedifferentiating cardiomyocytes. Cardiomyocytes were isolated from adult mice via retrograde perfusion and cultured in laminin-coated 96 well plates. Lineage tracing mice, carrying the Tamoxifen inducible Cre, *Tg(Myh6-cre)* (32), and *Rosa26^CAG-LSL-tdTomato^* reporter alleles (**Fig. 1a**), were used to permanently mark cardiomyocytes in culture, enabling unambiguous identification despite morphological and/or transcriptional changes during dedifferentiation. We found that cardiomyocytes isolated under these conditions can be cultured long term with high survival (**Fig. 1b**) (>50% after one week) and form networks that beat spontaneously and coordinately (Video S1). Morphological dedifferentiation occurs during the first 3-5 days of culture, as the cells adjust to the 2-dimensional substrate by rounding, probably due to the absence of axial mechanical stimulation (**Fig. 1b**). The cardiomyocytes continue to adapt during the first couple weeks of culture, as they form new connections with other cardiomyocytes and reorganize their sarcomeres (**Fig. 1b**). Transduction by an adenovirus vector carrying a GFP reporter showed strong gene expression after 3 days (**Fig. S1**). Furthermore, similar to in vivo studies (4,5), we found that adult mouse cardiomyocytes cultured in these conditions do not exhibit observable cell cycle activity (**Fig. 2**). Thus, this culture system is useful to screen for induction of proliferation by candidate genes using adenoviral vectors.

**Figure 1.**
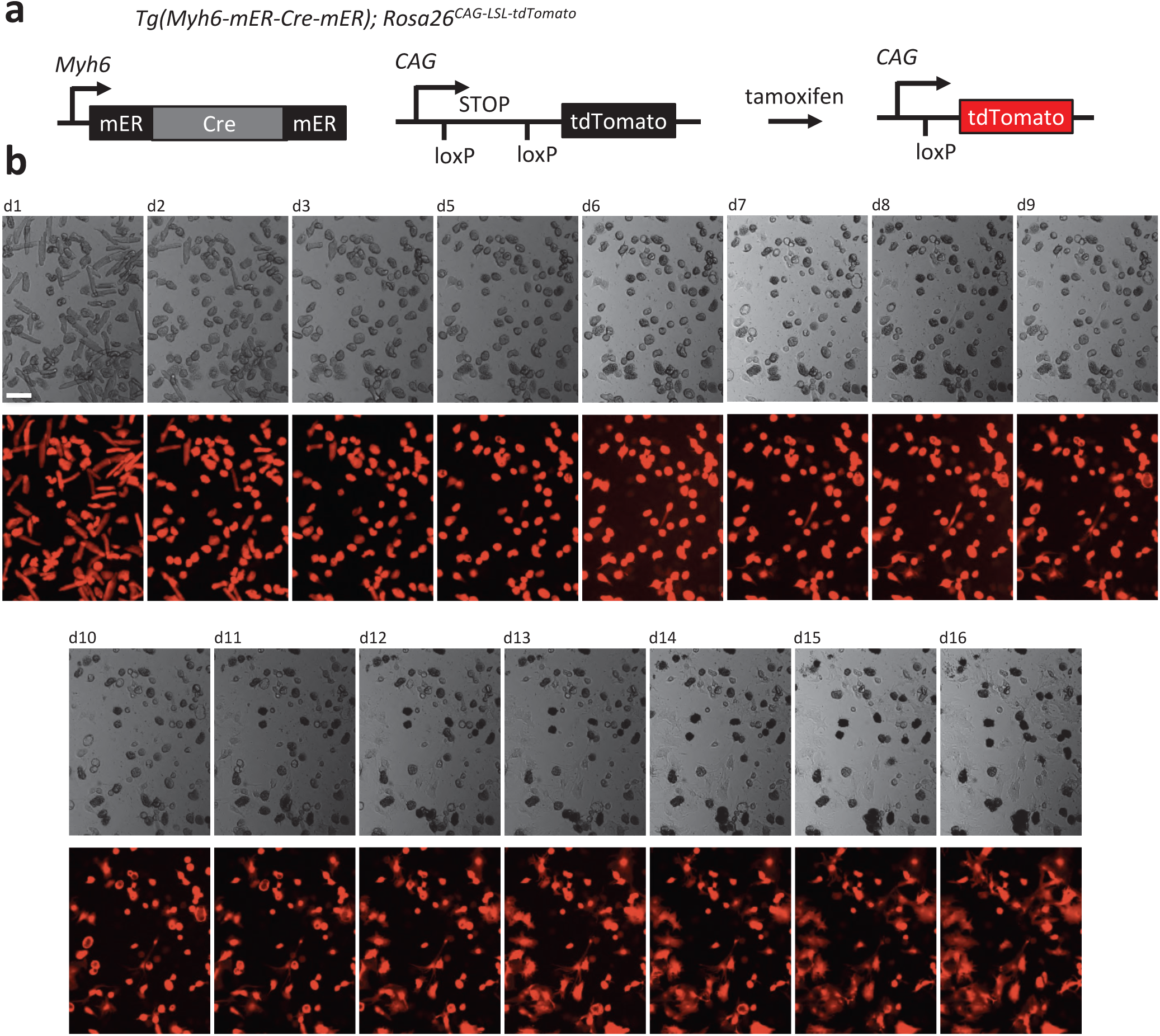
Live-cell imaging of genetically labeled adult mouse cardiomyocytes in culture. (a) Lineage-tracing transgenic mouse line was used to isolate adult cardiomyocytes, enabling unambiguous real-time identification during dedifferentiation. (b) Morphological changes of adult cardiomyocytes during dedifferentiation *in vitro* from day 1 (d1) to day 16 (d16). Cardiomyocytes are genetically marked by tdTomato before isolation and observed under the bright-field (top row) and fluorescent (bottom row) microscopy. Images of the same field are presented. Scale bar, 100 μm.

**Figure 2.**
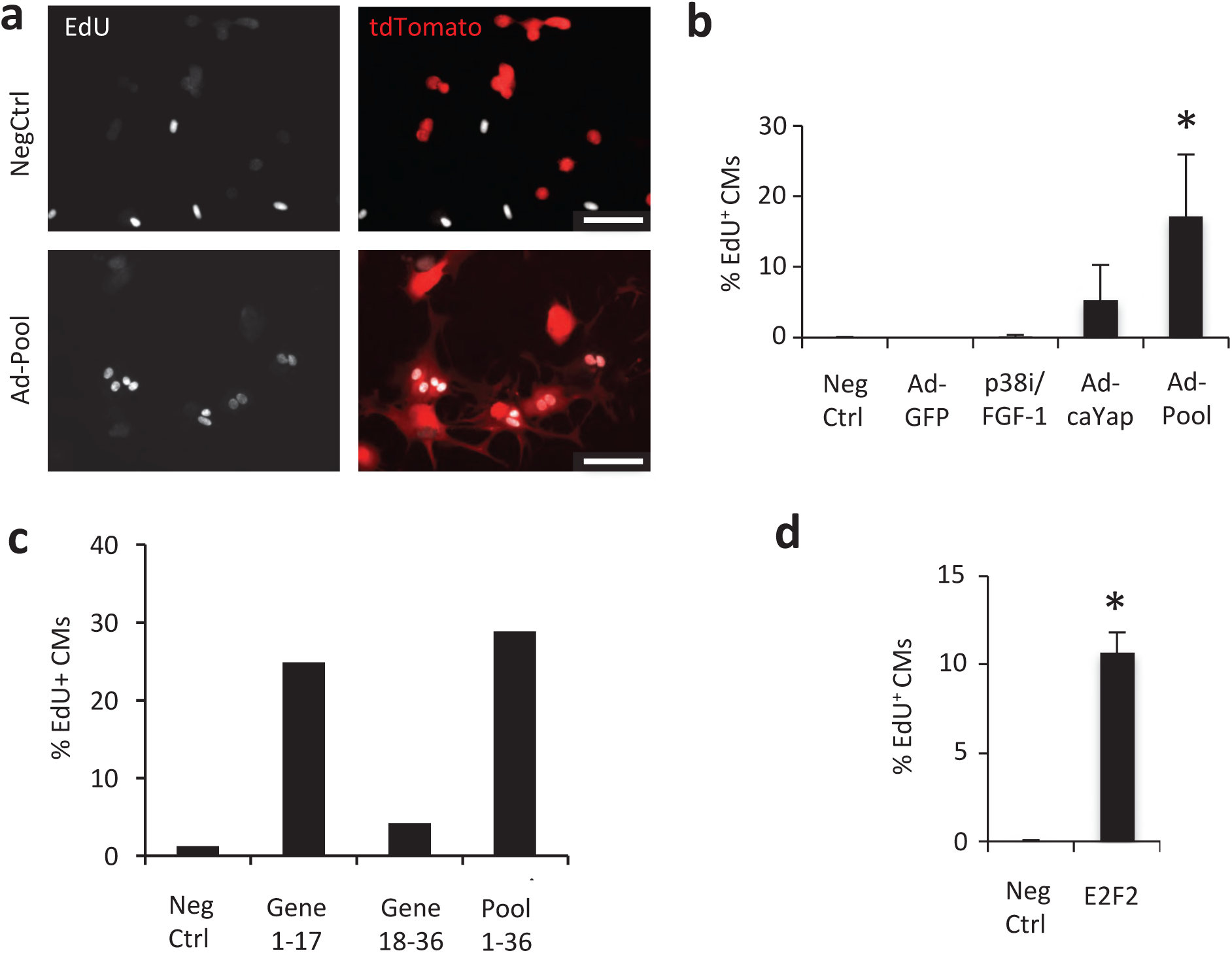
Candidate gene pool induces S-phase re-entry in cultured adult mouse cardiomyocytes. (a) Lineage-traced cardiomyocytes transduced with a pool of candidate genes show S-phase activity via EdU labeling of in situ fixed cardiomyocytes. (b) Quantification of S-phase induction by pooled candidate genes (1-36) compared to known cardiomyocyte cell cycle regulators p38i/FGF-1 and constitutively active Yap. Ad-GFP was used as a control for Adenovirus treatment. Negative control is no virus. (c) A subpool comprising genes 1-17 contains a candidate gene that is capable of activation of EdU incorporation in adult cardiomyocytes. (d) Identification of E2F2 that is sufficient to induce adult cardiomyocyte S-phase entry. Error bars represent standard deviation of mean from greater than or equal to 3 independent experiments. Asterisk indicates significance *p* < 0.05 compared to negative control. Scale bars, 100 μm.

### Candidate Gene Selection

It is well established that adult mammalian cardiomyocytes are highly resistant to cell cycle reactivation in physiological or pathological conditions. In contrast, the layer of epithelial cells that encapsulate the heart, or known as epicardial cells, are capable of cell division and rapid expansion in response to myocardial injury (33). Thus, we reasoned that master regulators of a proliferation-competent phenotype would still be active in epicardial cells, allowing them to re-enter the cell cycle in response to regenerative signaling. However, epicardial genes that are downregulated in adulthood are less likely to be master regulators required for cell cycle re-entry, and more likely are response genes that begin transcription downstream of activated master regulators.

Through gene expression comparison of epicardial and myocardial cells in both embryonic and adult hearts, we identified a list of candidate genes that are preferentially downregulated in the adult myocardium but not the adult epicardium (**Table S1**). Since our initial candidate genes contained novel factors with paralogs not represented by the microarray, such as those belonging to Bex/Tceal superfamily, we supplemented the gene pool with these additional genes. We also included 10 candidates based on reported functions in cardiomyocyte cell-cycle regulation, yielding a candidate gene list of 52 genes (**Table S2**). Due to low titers and difficulty cloning, some genes were not included in the final pool of 36 candidate genes that were functionally tested (**Table S3**).

### Candidate gene pool induces S-phase re-entry in adult mouse cardiomyocytes

Candidate gene ORFs were packaged individually into replication-defective serotype 5 adenoviral vectors (Ad5). To determine if there exists a combination of genes in the candidate gene pool that can induce adult mouse cardiomyocyte proliferation, we combined all candidate gene vectors into a single pool and delivered to cardiomyocytes en masse at day 1 post-isolation. S-phase re-entry was determined by labeling cardiomyocytes with EdU between days 2 and 7 post-transduction. Staining revealed strong cell cycle induction (∼17% EdU^+^ cardiomyocytes) from the pooled candidate genes (**Fig. 2 a, b**). We compared the effect of expressing the candidate gene pool to treatments that have been previously reported to stimulate cardiomyocyte proliferation. In our hands, combined treatment with FGF-1 and p38 inhibitor did not result in strong EdU labeling of adult mouse cardiomyocytes (**Fig. 2b**). Constitutively active Yap^S112A^ (caYap) did induce cell cycle re-entry in adult cardiomyocytes, consistent with previous reports (10,33) A control adenoviral vector expressing GFP, similar to the untreated control, did not exhibit a significant effect on proliferative activity. In addition, staining of a cell cycle marker Ki67 showed a similar result with ∼11% Ki67^+^ cardiomyocytes in the group transduced with the pooled viral vectors (**Fig. S2**).

### Narrowing candidate genes to those sufficient for S-phase re-entry

To identify candidate genes sufficient for S-phase re-entry, we applied subtractive pools to the cardiomyocytes and labeled with EdU as before. First the total gene pool was divided, in no particular order, into two subpools, genes 1-17 and genes 18-36 (**Table S3**). Delivery of these complementary subpools revealed subpool of genes 1-19, was significantly higher than the subpool of genes 18-36 pool for cell cycle induction (**Fig. 2a**), suggesting this subpool contains one or more candidate genes with strong cell cycle activity. This pool was further analyzed until a single virus transgene, that overexpresses E2f2, was found to be sufficient to drive adult mouse cardiomyocyte EdU incorporation (**Fig. 2d**). Interestingly, E2f2 is down-regulated only by ∼2 fold from E11.5 to adult epicardial cells but drastically by more than 20 fold from the E11.5 to adult myocardium (**Table S2)**.

### E2f2 is sufficient for S-phase re-entry, but induces cell death

E2f2 was previously reported to induce S-phase re-entry in cardiomyocytes (31), thus we hypothesized this gene may be partially responsible for the cell cycle activity of the candidate gene pool. However, in contrast to previous reports (30,31) we found that E2f2 overexpression, in the absence of other candidate genes, induced massive cell death in adult mouse cardiomyocytes under our culture conditions (**Figs. 3**, **S2**). Both total cardiomyocyte counting (**Fig. 3a**) and longitudinal tracking (**Fig. 3b**) revealed loss of more than 75% cardiomyocytes when E2f2 is overexpressed. TUNEL staining was positive in many cells at 5 days post-transduction (**Fig. S3**). However, we did not observe TUNEL positive cardiomyocytes that were still positive for the tdTomato lineage marker. Due to the small percentage of non-cardiomyocytes, it was unlikely that all of the TUNEL positive cells are not cardiomyocytes. Thus, in E2f2-induced cardiomyocyte death, it seems that genomic degradation, detected by the TUNEL assay, likely occurred after cell membrane disruption.

**Figure 3.**
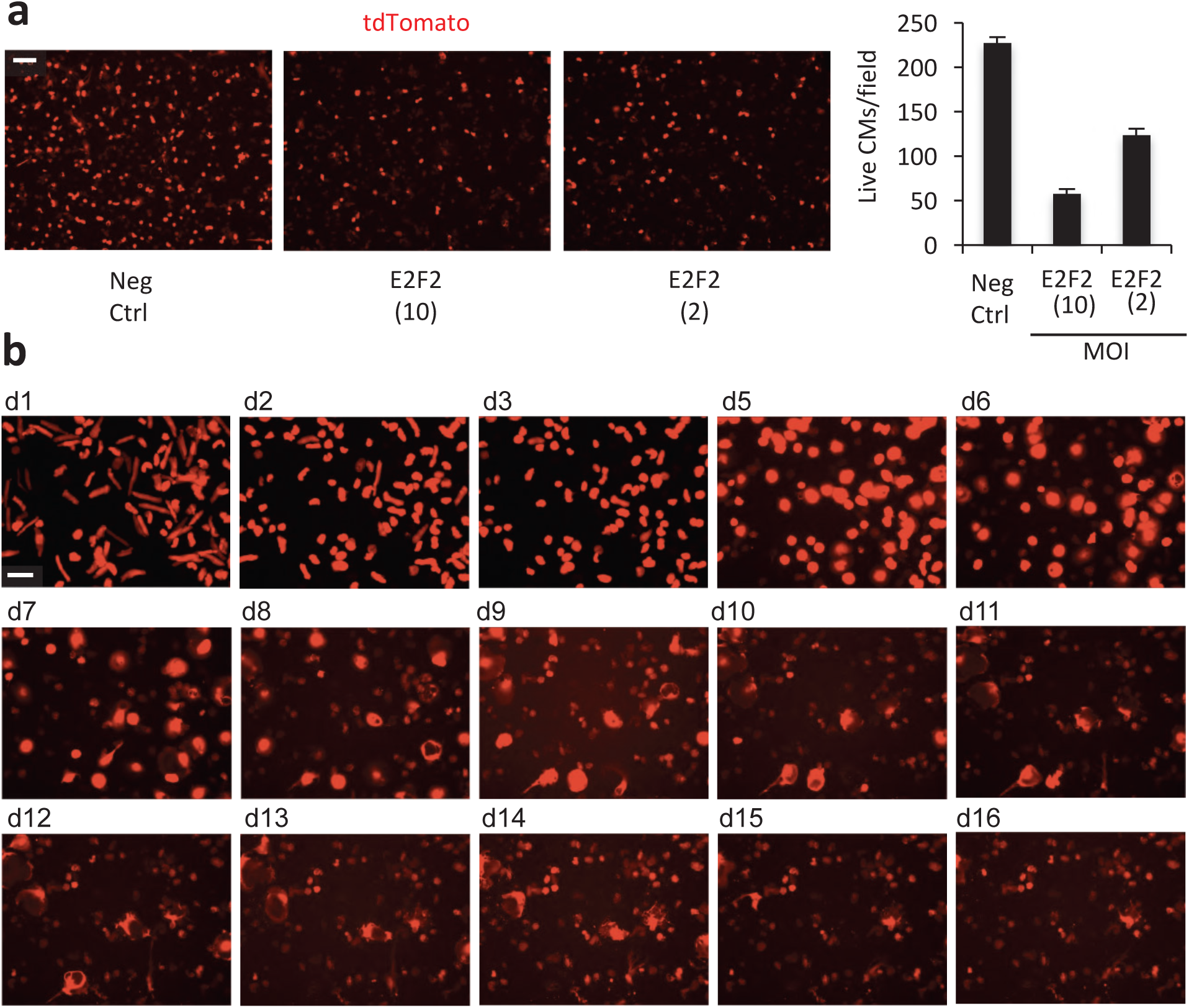
E2F2 overexpression induces adult mouse cardiomyocyte (CM) cell death. (a) E2F2 was expressed by adenovirus vector at various MOI. Cell viability is visualized by the transgenic tdTomato expression (red) at day 7 post-transduction and quantified on the right. (b) Longitudinal imaging of CMs revealed cell death in the E2F2-expressing group. Scale bar, 100 μm. Error bars represent standard deviation from duplicates in a single experiment.

### Tceal8 and BEX1 rescue cell death

Since E2f2 did not appear to cause cell death when co-expressed with the rest of the candidate genes in the virus pool (**Fig. S4**), we performed an additional screen to identify factors that can preserve cell viability in the presence of E2f2 overexpression. To that end, we noticed that Subpool 1.2 caused a similar degree of cardiomyocyte cell death to E2f2 alone (**Fig. S4**). Accordingly, we reasoned that one of the genes removed from Subpool 1.2 could rescue viability in the presence of E2f2. This Subpool lacked Tceal8, GOLM1, BMP4 and TUBB2B (**Fig. S4**). To test this, we co-expressed E2f2 with either Tceal8, GOLM1, BMP4, or TUBB2B, and then assessed cardiomyocyte viability at 7 days post-transduction (**Fig. S5**). Of these factors, only Tceal8 could rescue cell viability in the presence of E2f2 (**Fig. 4a and S5**). TUNEL staining confirmed a striking reduction of degraded genomic DNA (**Fig. S6**).

**Figure 4.**
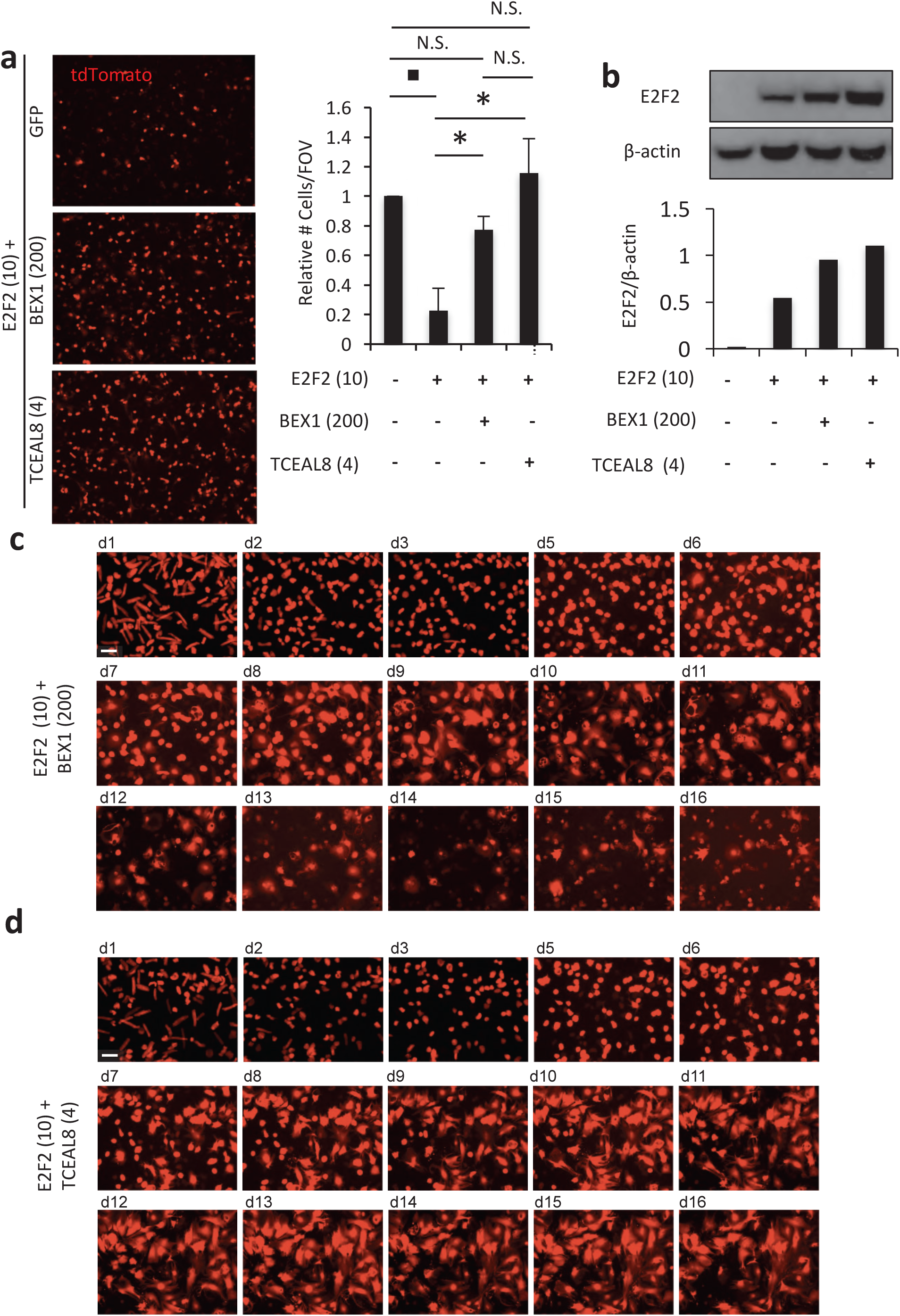
BEX1 and TCEAL8 delays or blocks E2F2-induced cell death in cultured adult mouse cardiomyocytes (CMs). (a) Cell viability is visualized by the transgenic tdTomato expression (red) at day 7 post-transduction, and quantification plotted at right. (b) Western blot (top) and densitometric analysis (bottom) of ectopic V5-tagged E2F2 expression using an anti-V5 antibody. (c, d) Longitudinal imaging of tdTomato-labeled CMs when E2F2 was co-expressed with BEX1 (c) or TCEAL8 (d). Scale bars, 100 μm. MOI is indicated in parentheses. \**p* < 0.05, N.S. = non-significant. ■ Indicates 95% confidence interval does not include 1. Error bars represent standard deviation from 3 independent experiments.

Since Tceal8 was effective at inhibiting cell death when co-expressed with E2f2, we suspected that other Bex/Tceal superfamily members may be able to regulate cell death in the context of E2f2 overexpression. To that end, we were able to clone other candidate genes belonging to the Bex/Tceal superfamily, Tceal5, Bex1, Bex3, and Bex4, and each was co-expressed with E2f2 (**Fig. S5**). Of these genes, Bex1 exhibited the strongest capacity to rescue E2f2-mediated cell death (**Fig. 4a and S5**). Western blot analysis of E2f2 showed that Tceal8 and BEX1 co-expression did not simply reduce E2f2 protein expression, but rather caused an increase in ectopic E2f2 protein accumulation (**Fig. 4b**).

Longitudinal imaging revealed that most E2f2-treated cardiomyocytes died between days 4 and 6 post-transduction (**Fig. 3b**). By contrast, when co-expressed with BEX1, most E2f2-treated cardiomyocytes survived through day 8 post-transduction (**Fig. 4c**). Interestingly, co-expression with Tceal8 prevented cell death in E2f2-treated cardiomyocytes through at least 14 days post-transduction (**Fig. 4d**).

### Tceal8 suppresses but Bex1 permits E2f2-induced cell cycle re-entry

Surprisingly, assessment of DNA synthesis with EdU labeling showed that Tceal8 strongly inhibited E2f2-induced cell cycle re-entry (**Fig. 5a**). However, in contrast to Tceal8, BEX1 facilitated strong cell cycle re-entry when co-expressed with E2f2 in adult mouse cardiomyocytes (**Fig. 5a**). This proliferative activity was also verified by Ki67 staining: E2f2 and BEX1 co-expression increases Ki67-positive cardiomyocytes to 27% (**Fig. S7**). We confirmed comparable expression levels of Tceal8 and BEX1 in our culture (**Fig. S8**). Altogether, these results suggest that Tceal8 and BEX1 have distinct functions in regulating E2f2-mediated gene transcription.

**Figure 5.**
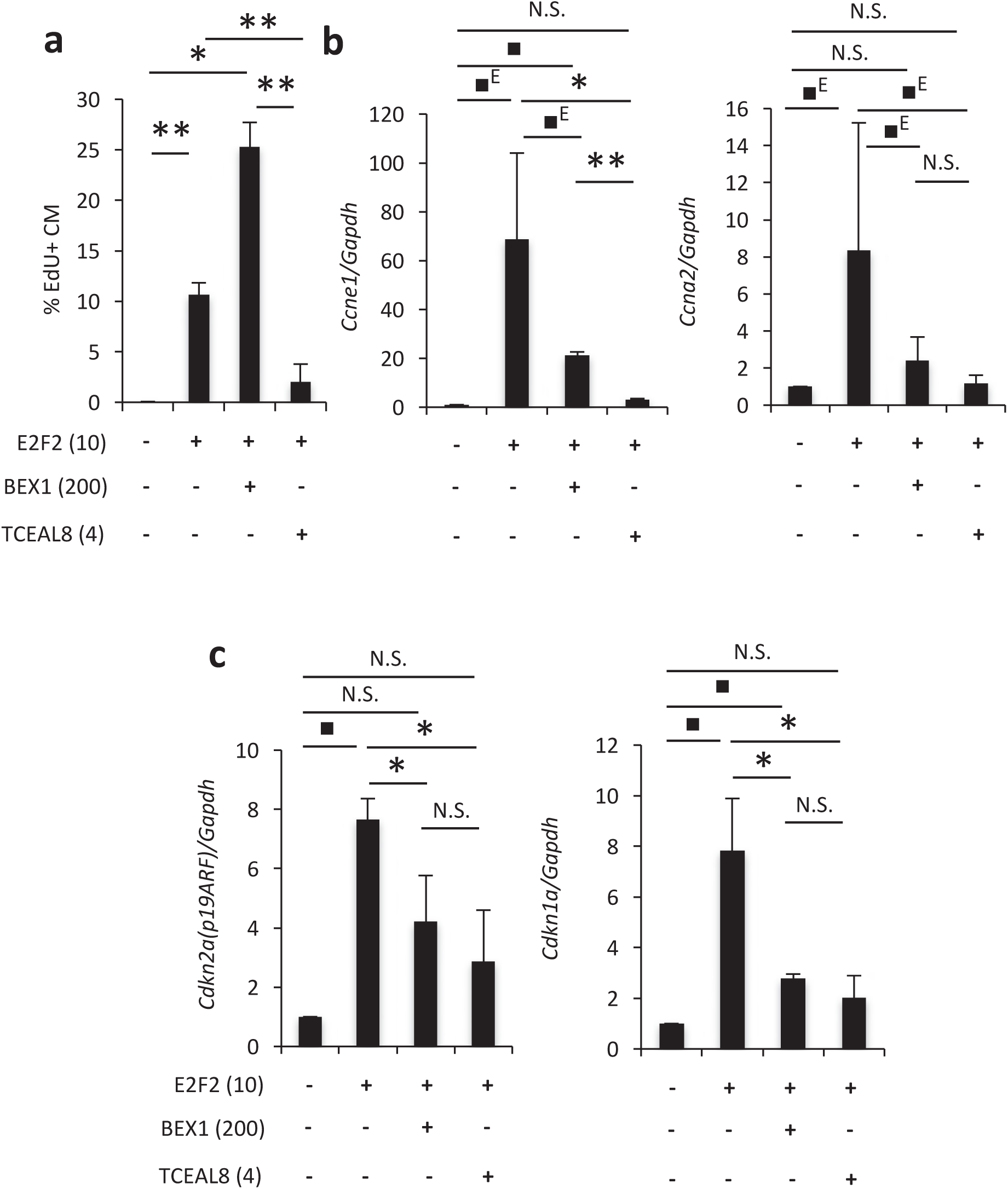
BEX1 and TCEAL8 inhibit cell death while modulating S-phase re-entry. (a) BEX1 permits S-phase re-entry by E2F2, but TCEAL8 co-expression strongly inhibits DNA synthesis. (b) RT-PCR analysis of positive cell cycle regulator genes *Ccne1* (Cyclin E) and *Ccna2* (Cyclin A2). (c) RT-PCR analysis of cell apoptosis regulator genes *Cdkn2a* (p19ARF) and *Cdkn1a* (p21CIP). Error bars represent standard deviation of mean from three independent experiments. \**p* < 0.05, \*\**p*<0.01, 95% confidence interval of mean does not include 1 when normalized to negative control (■) or E2F2(■^E^) N.S. = non-significant.

Real-time PCR showed that expression of endogenous positive cell cycle regulators Cyclin A2 and Cyclin E were induced by ectopic E2f2 expression, in agreement with\ its induction of S-phase (**Fig. 5b**). Tceal8 co-expression downregulated both Cyclin A2 and Cyclin E at the mRNA level, explaining the inhibitory effects of Tceal8 on cell cycle progression. In consonance with its permissive effect on DNA synthesis, BEX1 did not completely inhibit E2f2-mediated CyclinA2 and Cyclin E expression. Collectively, these data suggest that Tceal8 may block cell cycle activity in part by inhibiting Cyclin A2 and Cyclin E expression.

### BEX1 and Tceal8 expression is developmentally regulated in the heart

The observation that BEX1 rescues cell death while allowing cell cycle re-entry with E2f2 over-expression in adult cardiomyocytes is interesting and possibly relevant to the endogenous regulation of the cardiomyocyte cell cycle and maturation. Since murine cardiomyocytes are known to proliferate rapidly during early embryonic development, but permanently exit the cell cycle shortly after birth, the mapping of Bex1 expression with respect to these cell cycle and maturation dynamics could lead to a better understanding of Bex1 function. Thus, to help understand the potential function of Bex1 in vivo, we searched available databases (35) for the temporal dynamics of Bex1 expression in the developing heart. RNA-seq analyses showed high expression levels of Bex1 in the heart at E10.5, but a rapid reduction after birth (**Fig. 6a**). Interestingly, Tceal8 gene expression follows the same temporal trend (**Fig. 6b**). The drastic decreases of Bex1 and Tceal8 expression in the neonatal heart suggest that they may play important functions in cardiomyocyte cell-cycle exit and maturation *in vivo*.

**Figure 6.**
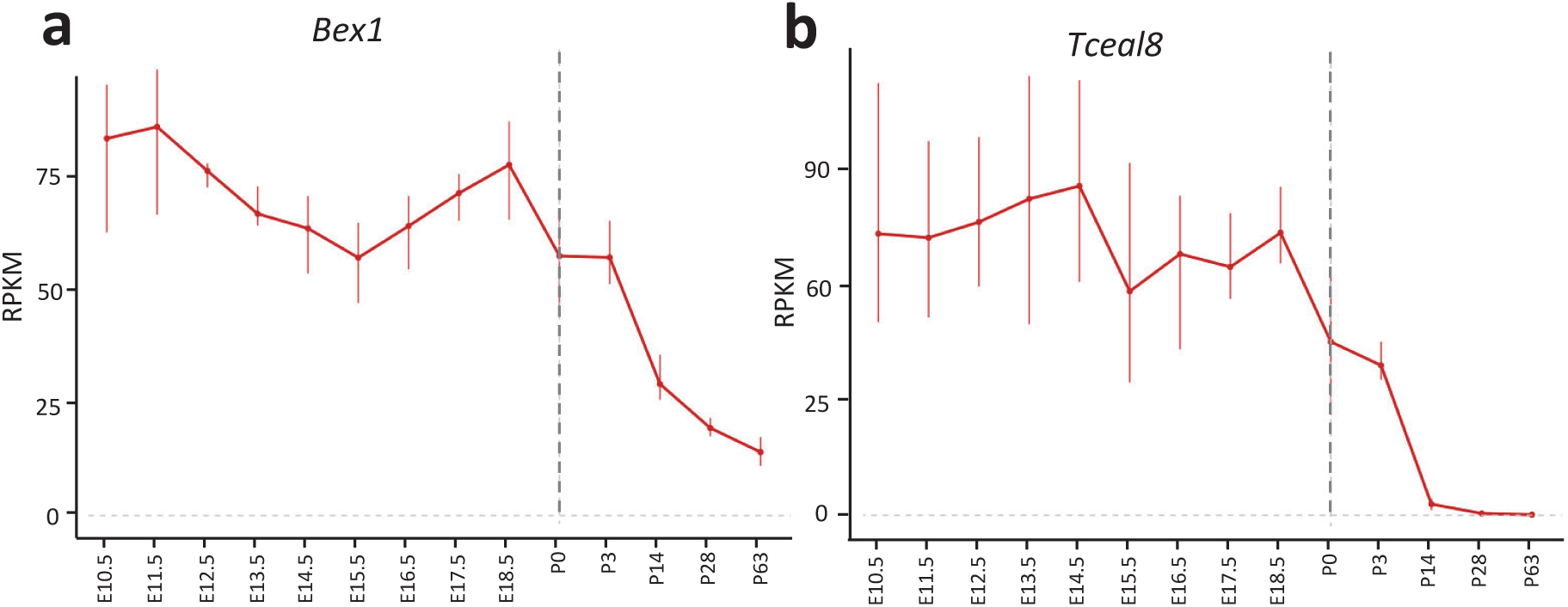
*Bex1* and *Tceal8* expression in mouse heart development. Data are derived from published RNA-seq data. Error bars represent standard deviation.

## DISCUSSION

Several reports have demonstrated increased cell cycle activity by ectopic gene expression in proliferative neonatal mammalian cardiomyocytes (36–38), but fewer have shown cell cycle re-entry in senescent adult mammalian cardiomyocytes. Of factors known to be sufficient for induction of DNA synthesis in adult mammalian cardiomyocytes, strong evidence exists for E2F transcription factors (31,39), thus they are of interest to the pressing question of how adult mouse cardiomyocytes can be artificially stimulated to proliferate (40,41). E2F transcription factors 1-3 are known activators of the cell cycle and are negatively regulated through sequestration by pocket proteins, such as retinoblastoma protein (RB), p107, and p130 (42). Overexpression of E2F2 here by the CMV promoter likely overwhelms pocket protein sequestration of E2Fs, thus resulting in high levels of the activated free form E2F2, which upregulated Cyclin expression and DNA synthesis (**Fig. 5**) (30,31).

Long-term culture did not result in cytokinesis, rather all cells treated with Ad-E2f2 eventually died. The exact type of cell death is unclear, but may be related to several potentially overlapping and concurrent death processes (43). Notably, we observed positive TUNEL staining in Ad-E2f2-treated cardiomyocytes, but did not observe TUNEL positive cells that were also positive for the lineage tracing marker TO. Previous work showed that fluorescent protein expression is diminished in apoptotic cells, but that necrotic cells almost completely lose fluorescent signal (44). Thus, it appears that genomic degradation may have occurred after membrane permeabilization due to forms of cell death other than apoptosis, such as necrosis (45–47). It should be noted that Bex1 has been shown to inhibit apoptosis in the rat central nervous system (48). Although our data suggest a cellular death mechanism other than apoptosis (Fig S3), there may be common pathways involved in multiple forms of cell death (49).

The primary cause of cell death could potentially be through dysregulation of E2f2 temporal dynamics or expression levels, leading to death signaling due to failed cell cycle checkpoints. In post-mitotic neurons, E2F activation can lead to cell cycle re-entry followed by cell death (50). This phenomenon seems to be cell context dependent and may play a role in neurodegeneration (51). Furthermore, E2f2 and E2F1 were reported to provide genomic stability in some neurons in the context of DNA damage (52). This study additionally showed accumulation of E2f2 protein levels, despite unchanged transcription. This is especially interesting considering our observation that BEX1 and Tceal8 appear to also allow the accumulation of E2f2 protein, detected via western blot (Fig. 4b). Additionally, since E2f1 is also sequestered by Rb, it is possible that a significant proportion of endogenous E2f1 is available in its activated form under these conditions, potentially contributing to observed cell death. Finally, during the first several days of culture, when gene delivery is taking place, the cells are only loosely attached. Thus, it is possible that delivery of viruses to CMs later in culture when they have adapted to culture conditions and are well-anchored may have a different effect. In future work, it would be interesting to test the effect of drug-inducible E2f2 expression at different time points in order to see if transient E2f2 expression can enable post-mitotic cardiomyocyte cell cycle re-entry without cell death.

The mechanism of rescue from cell death and cell cycle modulation by Bex1 and Tceal8 are unclear, but may be suggested by RT-PCR data (Fig. 5). Specifically, p21CIP1 (53,54) and p19ARF (55,56) are involved in cell cycle control and cell death decisions, and have even been reported to interact (57,58). For example, p19-ARF, in addition to other functions, appears to induce apoptosis in response to E2F overexpression (59). Since our data suggest non-apoptotic cell death, it is further interesting that a short form of p19ARF was reported to contribute to caspase-independent cell death (60). P21, like p19, has also been shown to have multiple roles in coordinating the cell cycle with cell death decisions, and is additionally involved in thryoid-mediated cardiomyocyte maturation (61). Moreover, p21 has been shown to oppose E2F signaling, which may explain why its expression was increased in response to damaging levels of E2f2 overexpression (62). Since p19ARF is known to target E2F transcription factors for proteasomal degradation (50), it is also possible that accumulation of E2F2 by Bex1 and Tceal8 could be through suppression of p19ARF (Fig. 5). Furthermore, downregulation of p21 and p19, both having roles as cell cycle inhibitors, could potentially explain how BEX1 facilitates S-phase entry. Why Tceal8 does not facilitate S-phase re-entry under our conditions, despite downregulation of both p21 and p19, could be simply related to its stronger effect on Cyclin E downregulation. Furthermore, Tceal8 appeared to allow S-phase re-entry when co-delivered with Subpool 1, but not when only co-delivered with E2f2 (**Figs. 2a, 5a**); thus, there may be other factors in Subpool 1 responsible for the discrepancy.

The endogenous function of Bex/Tceal family proteins are largely unknown, but previous work has shown their role in other cell types such as neuronal differentiation (63) and skeletal myoblast maturation (64,65). BEX1 has been shown to regulate NGF signaling in neurons by altering its activity on p75^NTR^ and trkA. The alteration in NGF signaling by BEX1 prevents downstream activation of NFkB and thereby prevents neuron differentiation. Furthermore, NGF has a prosurvival effect on rat cardiomyocytes (66) and long-term NGF signaling may contribute to myocardial deterioration in chronic heart failure (67). Here, it is tempting to speculate that Bex1 (and possibly Tceal8) may be involved in transducing death and differentiation signals in cardiomyocytes through p75^NTR^ in concert with SC-1, as has been previously suggested (68). In support of this hypothesis, SC-1 has been shown to inhibit cell cycle progression by repressing Cyclin E transcription (69). A possible role for Bex1 here would be consistent with our observation that Bex1 (or Tceal8) overexpression downregulates Cyclin E expression induced by E2f2. Thus, the p75^NTR^/SC-1 signaling axis may provide a mechanism for the ability of Bex1 to inhibit cell death. On the other hand, since our experiments relied solely upon Adenovirus-mediated gene delivery, it cannot be ruled out the possibility that phenotypes observed in this work are affected by virus-mediated signaling.

Since Bex1 has been shown to interact with many different proteins, several potential mechanisms, independent of p75^NTR^ signaling, are suggested by previous work. For example, BEX1 has been reported to bind Calmodulin (65), BAG4/SODD, and Caspase 8 (70), all of which have roles in death signaling through DISC (71–73). In contrast to our observations, Bex1 has been shown to have a pro-apoptotic role by binding to Bcl2 in cancer cells (74). However, it is unclear what role the cellular context plays in determining the downstream outcome of this interaction. Bex family proteins have also been shown (70,75) to interact with several common proteins - RAD51, ATM, AKT, AURKA, c-JUN, CHEK2 – as does BRCA1, which regulates DNA damage through the BASC complex (76), and is required for CCNE1 amplified tumors (77). Alternatively, Bex1 interactor Pax2 (70) has been shown to regulate retinoblastoma protein (78), a particularly interesting observation given the role of RB in controlling E2F2 activity. Moreover, BEX1 binds FBXW7 (70), an F-box protein which promotes Cyclin E degradation through the SKP1-Cullin1-F-BOX (SCF) E3 ligase complex (79). This interaction could help explain how Bex1 (and possibly Tceal8) reduces Cyclin E expression in response to ectopic E2f2. Furthermore, high Cyclin E levels have been shown to selectively degrade activator E2Fs (80), potentially explaining why co-expression of Bex1 or Tceal8 actually increases E2f2 protein levels in contrast to Cyclin E suppression (**Figs. 4,5**). Finally, PTEN, may interact with Bex1, as well as several common interacting proteins, such as AKT, CASP8, FBXW7, STK11, FZR1, and MDFI (81), and PTEN functionally interacts with p19ARF (82), p21CIP (83), and regulates cyclins (84,85). Moreover, PTEN has been shown to be an effector of apoptosis inhibition by retanoic acid (86). Thus, PTEN should be considered as a potential molecular player in Bex1 signaling in the context of cardiomyocyte maturation and survival.

In recent work, Bex1 was found to mediate pro-inflammatory signaling by selective modulation of AU-rich mRNA targets in a model of heart failure (87). Thus, it is conceivable that Bex family-mediated RNA processing could play a role in shaping the cardiomyocyte response to E2f2-induced stress, but further investigation would be required. RNAseq data in rats from those experiments, deposited by Accornero et al, in contrast to our results, show that Bex1 overexpression actually increased Cyclin and cyclin-dependent kinase inhibitor (cdki) expression, including Cyclin E and p21CIP. Thus, it is unclear how the cellular context affects the action of Bex1 on cell cycle protein expression. For example, it is possible that Adenovirus signaling could play a role in shaping the response of CMs to both E2F2 and Bex1/Tceal8 overexpression. Alternatively, considering the mild phenotypes in Bex1 knockout animals, it is conceivable that Bex1, and possibly other Bex family proteins act to fine tune cyclin expression in order to modulate cell cycle and cell death decisions, thus explaining how Bex family proteins may be able to increase or decrease target transcript/protein expression depending on the cellular context.

Aside from the aforementioned potential mechanisms for Bex superfamily mediated regulation of cell death and proliferation, it is interesting to consider the Hippo pathway, due to its role in cardiac proliferation, regeneration, and cell death (10,19–23) as well as its regulation via WW domain interactions (88) and due to the interaction between Bex1 and the Hippo- and NFKB-interacting protein LMP1 (89). YAP, a major signaling protein in the Hippo pathway, is known to interact with proteins via a WW domain by binding to proline rich recognition sequences (90). Tceal9 (Wbp5) has been shown to induce Yap nuclear localization (91), a sign of Yap activation. Since TCEAL9 contains a WW binding domain and TCEAL8 contains a proline rich sequence at its C-terminus, it is possible that other Bex/Tceal superfamily members may interact with Yap and could possible account for the activity of Bex1 or Tceal8 in this work. Thus, we tested to see if Bex/Tceal superfamily members from our screen can activate Yap in neonatal cardiomyocytes. Of the genes tested, Tceal5 (which exhibited mild rescue of cell viability; **Fig. S5**) increased nuclear Yap (**Fig. S10**). Although Bex1 and Tceal8 did not increase nuclear Yap, it is possible that the cellular context could play a role in the outcome of Hippo signaling (92,93), as neonatal cardiomyocytes are quite different from adult cardiomyocytes (**Table S1**). Furthermore, miRNA signaling has been shown to control Tceal family proteins in the context of Yap signaling (91).

It should be noted that the specific MOIs used in this study could have played a role in the differential effect between Bex1 and Tceal8, but in our hands lower MOIs of Bex1 had no effect and higher MOIs caused cellular toxicity (data not shown). Lower MOIs of Tceal8 did not allow S-phase re-entry while rescuing cell death. Thus, it appears the actions of Bex1 and Tceal8 on S-phase re-entry are distinct. It is also worth mentioning that the ectopically expressed BEX1 used in this experiment is of human origin, while E2f2 and Tceal8 are derived from mouse cDNA. Thus, it is possible that mouse Bex1 and human Bex1 could have different effects. It is also possible that Bex1 derived from mouse could have a similar effect at a lower MOI. Expression in Ad293 cells showed similar protein levels between Bex1 and Tceal8 despite Ad-Bex1 being delivered at 40X higher MOI than Ad-Tceal8 (**Fig. S8**). This could be due to differential silencing of the Bex1 transcript, inefficient translation, or protein degradation, distinct from Tceal8. It is also possible that the difference in gene function of Tceal8 and Bex1 could have affected the virus titer assay.

The interplay between death and differentiation signaling is becoming increasingly appreciated (64,94–96). Our results showed that Bex1 is highly expressed in early stages of myocardial development and that Bex1 and Tceal8 inhibited cell death in response to ectopic E2f2. Furthermore, Bex family proteins have been suggested to behave as disordered signaling hubs (75), involving their interaction with many different protein partners. Collectively, these data and ideas suggest that Bex family proteins may act at several different levels to coordinate the balance between cellular decisions of death, division and differentiation during early stages of heart development and cardiomyocyte maturation.

## METHODS

### 1. RNA expression

#### 1.1 Microarray analysis for candidate gene selection

In order to compare gene expression between the epicardium and myocardium in both adult and embryonic hearts, six groups of RNA were prepared:

1. E9.5 *Tcf21^LacZ^* proepicardial organ (PEO) lacZ-positive cells
2. E9.5 *Tcf21^LacZ^* proepicardial organ (PEO) lacZ-negative cells.
3. E11.5 *Tcf21^LacZ^* heart lacZ-positive cells.
4. E11.5 *Tcf21^LacZ^* heart lacZ-negative cells.
5. Adult epicardium-enriched cells.
6. Adult hearts.

Tcf21 is a marker for embryonic epicardial cells (97), thus in *Tcf21^LacZ^* mouse embryos, epicardial cells express the lacZ transgene.

To isolate epicardial cells from E9.5 and E11.5 *Tcf21^LacZ^* mouse embryos, CD-1 females were mated with *Tcf21^LacZ^* males and hearts were harvested from 25-40 E9.5 or E11.5 embryos. Cells were dissociated by trysinization at 37°C for 20 minutes and resuspended in 100 μL prewarmed Leibovitz’s L15 medium (Invitrogen). Next, single cell suspensions were mixed with 100 μL prewarmed 2 mM fluorescein di-β-D-galactopyranoside (FDG) (Invitrogen) diluted in water and then incubated in a 37°C water bath for 1-2 minutes to permit substrate uptake. Finally, 800 μL ice-cold L15 medium was added to the mixture and the cells were placed on ice for 30 to 60 minutes before fluorescence activated cell sorting (FACS). Cells were sorted directly into Trizol solution for RNA extraction.

Because most adult epicardial cells don’t express *Tcf21* any more, adult epicardium-enriched RNA was isolated from the lysates after dissolving the surface (epicardial) cells of an entire adult ventricle from a 2-month old mouse in Trizol for 1 min. For adult heart, RNA was isolated from the entire adult ventricle from a 2-month old mouse.

RNA was extracted with the miRNeasy kit (Qiagen) following manufacturer’s instructions. For microarray analysis, 500 ng RNA and an Illumina mouse-6 gene chip were used.

Microarray data were obtained and analyzed by the Microarray Core facility of University of Texas Southwestern Medical Center.

#### 1.2 RNA expression in vitro

RNA extraction was performed on cardiomyocyte cultures by Trizol (Invitrogen) following manufacturer’s instructions. Briefly, culture dishes were placed on ice, media was removed and Trizol added, followed by homogenization by repeated pipetting. RNA was then extracted by ethanol-stabilized chloroform (Acros Organics), supplemented with 5 ug glycogen (Invitrogen) co-precipitant and precipitated in 0.3M sodium acetate, pH 5.5. Pellets were washed in 75% ethanol, followed by resuspension in RNAse-free water. cDNA was prepared using iScript Supermix (BIO-RAD) according to manufacturer’s instructions. qPCR was performed on 7900 HT thermal cycler (Applied Biosystems) in 386 well plates using SYBR Select Master Mix (Life Technologies). RT-PCR primers are listed in Table S4. Changes in gene expression, compared to control samples, were calculated using ΔΔCt relative quantitation (98).

### 2. Western Blot

Protein samples were obtained by replacing cell culture medium with 1% sodium dodecyl sulfate containing 1X protease inhibitor cocktail (Sigma P8340). Separation was performed in 4-12% Bis-Tris NuPAGE gels (Invitrogen) in MES-SDS buffer (Invitrogen) under reducing conditions. Proteins were then transferred to PVDF-Plus membranes (Spectrum Chemical) and blocked in 5% milk in TBST (0.1% Tween 20). Blots were probed overnight at 4 deg C with rabbit anti-V5 (Cell Signaling Technologies; 1:1000 in 5% BSA, 0.1% Tween 20, TBS) or for 1 hr at room temperature with mouse anti-β actin (GeneTex; 1:2000 in 1% BSA, 0.1% Tween 20, TBS), mouse anti-Bex1/2 or anti-E2f2 (Santa Cruz Biotech sc-376342 or sc-9967; 1:100 in 1% BSA, 0.1% Tween 20, TBS) primary antibodies. Anti-mouse or -rabbit secondary antibodies (GE) were incubated in 5% milk in TBST for 1 hour at room temperature. Western lightning enhanced chemiluminescence substrate was applied and blots exposed to Hyblot CL film (Denville). Semi-quantitation of western blots was done in Fiji by subtracting background with a 50 um radius, followed by integrated intensity measurement.

### 3. Adenovirus vectors

#### 3.1 Candidate gene cloning and virus production

Candidate gene ORFs were cloned into gateway entry vector pDONR-221 (Invitrogen) via PCR amplification with primers containing the attB recombination sequences. After sequence verification, entry vectors were recombined with the adenovirus type 5 destination vector pAd5-CMV-V5-DEST using LR Clonase (Invitrogen). pAd5 vectors carrying candidate gene vectors were further sequenced to confirm gene identity, and then digested with Pac I (NEB) for plasmid linearization. After phenol-chloroform extraction and ethanol precipitation, linearized plasmids were transfected into Ad293 producer cells by polyethyleneimine complexation. Adenoviruses were harvested when a 50% cytopathic effect was observed. Cells were collected by cell scraping in culture media, followed by centrifugation at 1000G x 5 min and concentration by removing all but 2 ml of media. Lysates were prepared by 4 freeze/thaws, cleared by centrifugation and stored at -80 degC.

#### 3.2 Virus Titer

Virus titer was determined by infecting Ad293 cells with serial dilutions of virus lysate. Cytopathic effect was scored and the dilution at which isolated viral plaques were observed was used to determine the concentration of infectious units (IU). Vectors exhibiting titers less than 1e5 IU/ml were considered low titer and were not evaluated. Each multiplicity of infection (MOI) reported in this work refers to the number of virus IU (determined in Ad293) per cardiomyocyte. Thus, the effective MOI in cardiomyocytes may differ depending on the relative permissivity between Ad293 and cardiomyocytes.

### 4. Cardiomyocyte isolation and culture

Cardiomyocytes were isolated by retrograde perfusion of collagenase and protease XIV to digest extracellular matrix as previously described (99). Cells were plated at approximately 15,000 cells/cm^2^ laminin (Corning) coated plasma-treated polystyrene in 96 well plates (Corning). Cells were cultured in Modified Eagle’s Medium (Corning) with 10 % fetal bovine serum (JRS) and Primocin antibiotic (Invitrogen). Half-change of media was performed every 2-4 days. Cardiomyocytes were typically transduced by adding virus cell lysate on day 1 post-isolation. For viability assay, live cells were counted manually, as determined by the presence of positive tdTomato fluorescent signal.

### 5. DNA synthesis

S-phase activity was detected by addition of EdU (Santa Cruz Biotechnology) to cell culture media between days 2-7 post-transduction. Staining was performed on formaldehyde fixed samples after permeabilization with 0.1% Triton X-100 and blocking with 3% bovine serum albumin (Santa Cruz Biotechnology) in phosphate-buffered saline. Briefly, staining solution (0.1 M ascorbic acid, 50 mM Tris-Cl, 1 mM CuSO4, pH 7) containing 10 uM Sulfo-Cyanine5 azide dye (Lumiprobe) was applied for 10-20 minutes at room temperature, washed in PBS and followed by Hoescht 33342 (Gibco) nuclear counterstaining and several PBS washes. EdU+ cells were counted manually in ImageJ using PointPicker.

### 6. Animal Use

All procedures and experiments have been approved by the Institutional Animal Use and Care Committee at the University of California, San Francisco. All experiments and procedures were performed in accordance with relevant guidelines and regulations.

### 7. Statistics

For all counts used in statistical tests, including EdU+ cells and cell number counts, >100 total cells per sample (except for cells treated with E2f2 alone) were counted in ImageJ using PointPicker. For cell culture samples treated with E2f2 alone (not in combination with Bex1 or Tceal8), the large degree of cell death resulted in lower cell counts (∼20-50 per sample). P-values were derived using one-sided Welch’s test in excel. For RT-PCR data normalized to negative controls (Fig. 5), statistical tests comparing to negative control were performed using a one-sided lower confidence interval for the mean of the treated group using the student’s t-distribution. RT-PCR data not comparing to negative controls were evaluated using the Welch’s test in excel.

## ACKNOWLEDGEMENTS

The present work is supported by NIH T32HL007731 (J. J.), NIH Pathway to Independence Award, Edward Mallinckrodt Jr. Foundation, March of Dimes Basil O’Conner Scholar Award, American Heart Association, American Federation for Aging Research, Life Sciences Research Foundation, Program for Breakthrough Biomedical Research, UCSF Eli and Edythe Broad Center of Regeneration Medicine and Stem Cell Research, Resource Allocation Program, and Cardiovascular Research Institute (G. N. H.).

## AUTHOR CONTRIBUTIONS

JJ performed experiments and wrote the manuscript. JL assisted with molecular cloning and virus generation. GH performed experiments and edited the manuscript.

---

**Figure S1.**
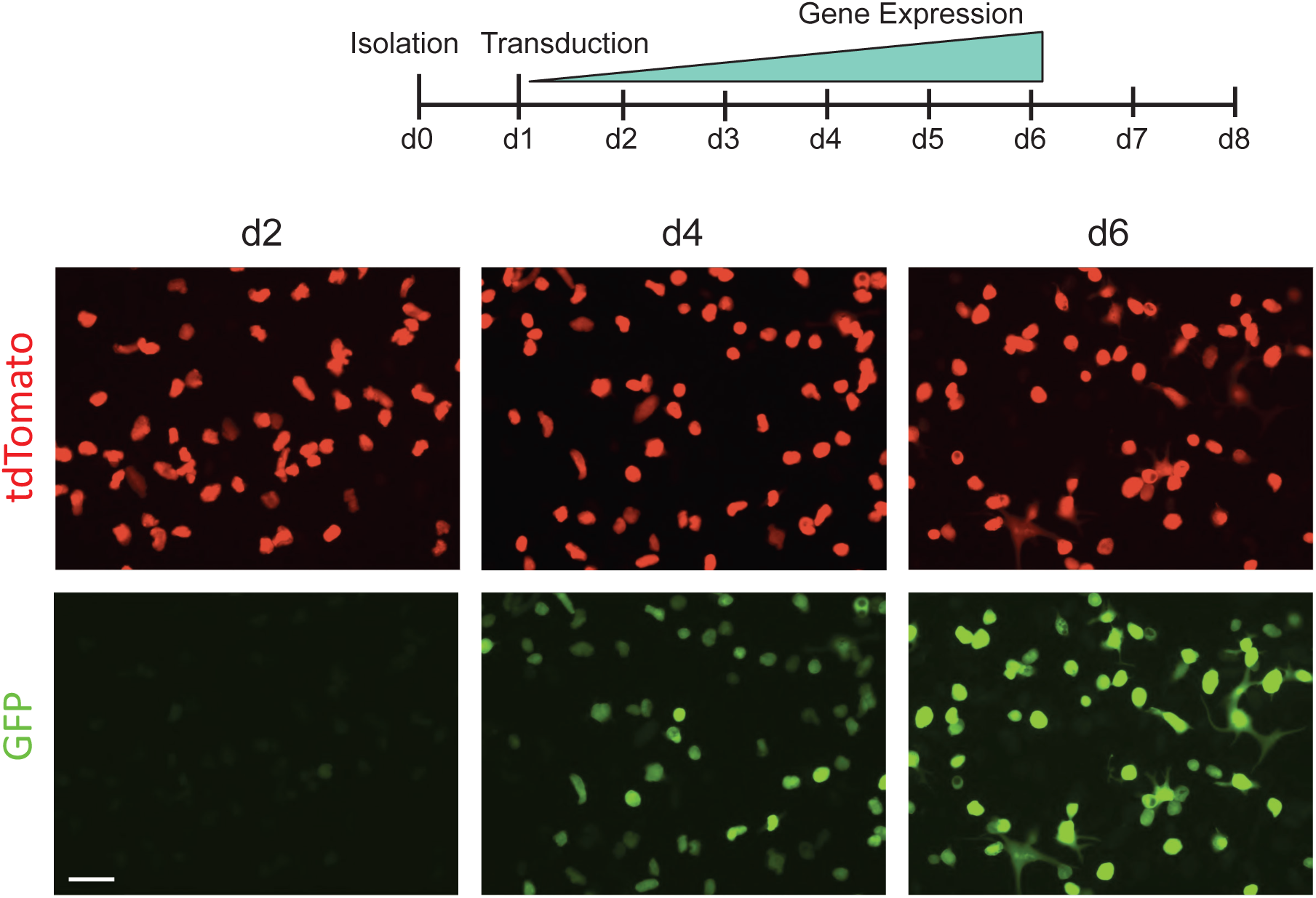
Transduction of cardiomyocytes by adenovirus expressing GFP. Lineage traced tdTomato^+^ adult mouse cardiomyocytes are transduced with Ad-GFP at day 1 post-isolation. Gene expression is detectable, but very low at d2 (d1 post-transduction), but robust at d4 and very strong by d6. Scale bars, 100 μm.

**Figure S2.**
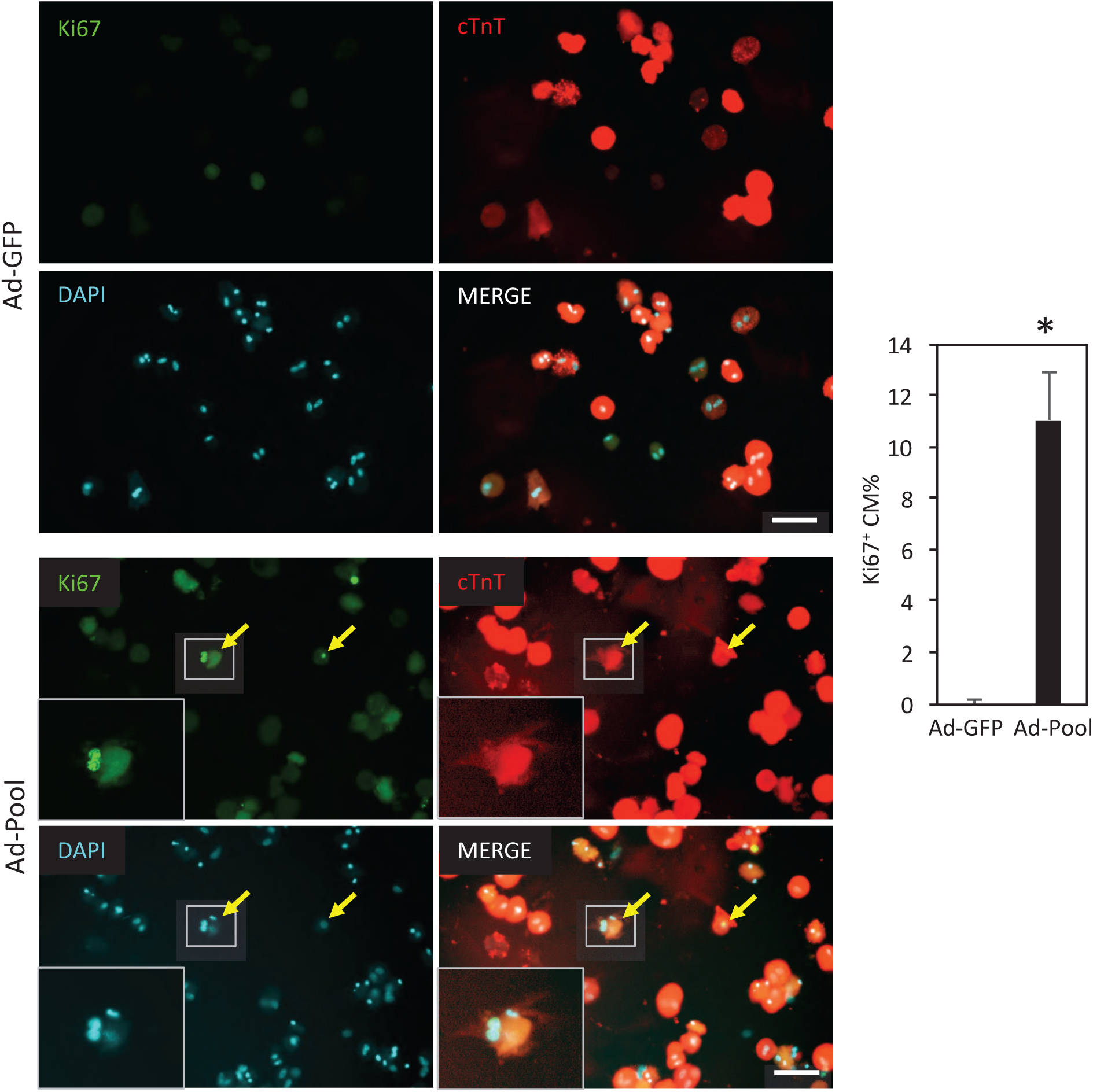
Increase of cardiomyocyte proliferation by pooled adenoviruses expressing all candidate genes. Adult mouse cardiomyocytes are transduced with viruses at day 1 post-isolation and stained at day 4. Yellow arrows point at Ki67-positive cardiomyocytes. High-magnification views of the boxed areas are shown in the insets. *, p<0.05. Error bars represent standard deviation of mean (n=3). Scale bars, 100 μm.

**Figure S3.**
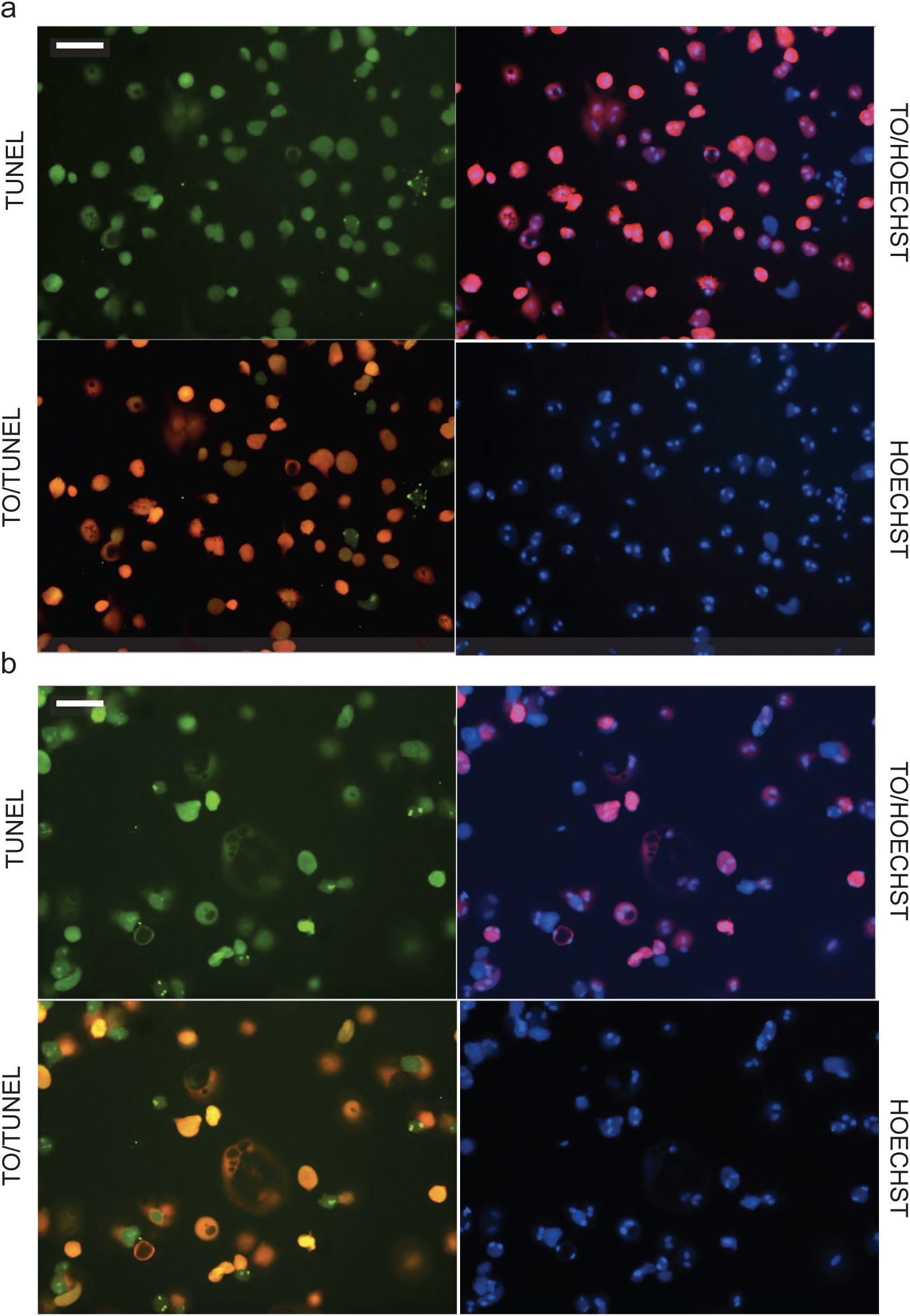
TUNEL staining of control and E2F2-treated cardiomyocytes. (a) TUNEL staining of untreated negative control cardiomyocytes shows low incidence of genomic degradation. (b) E2F2-treated cardiomyocytes by contrast show significant TUNEL label. Scale bars, 100 μm.

**Figure S4.**
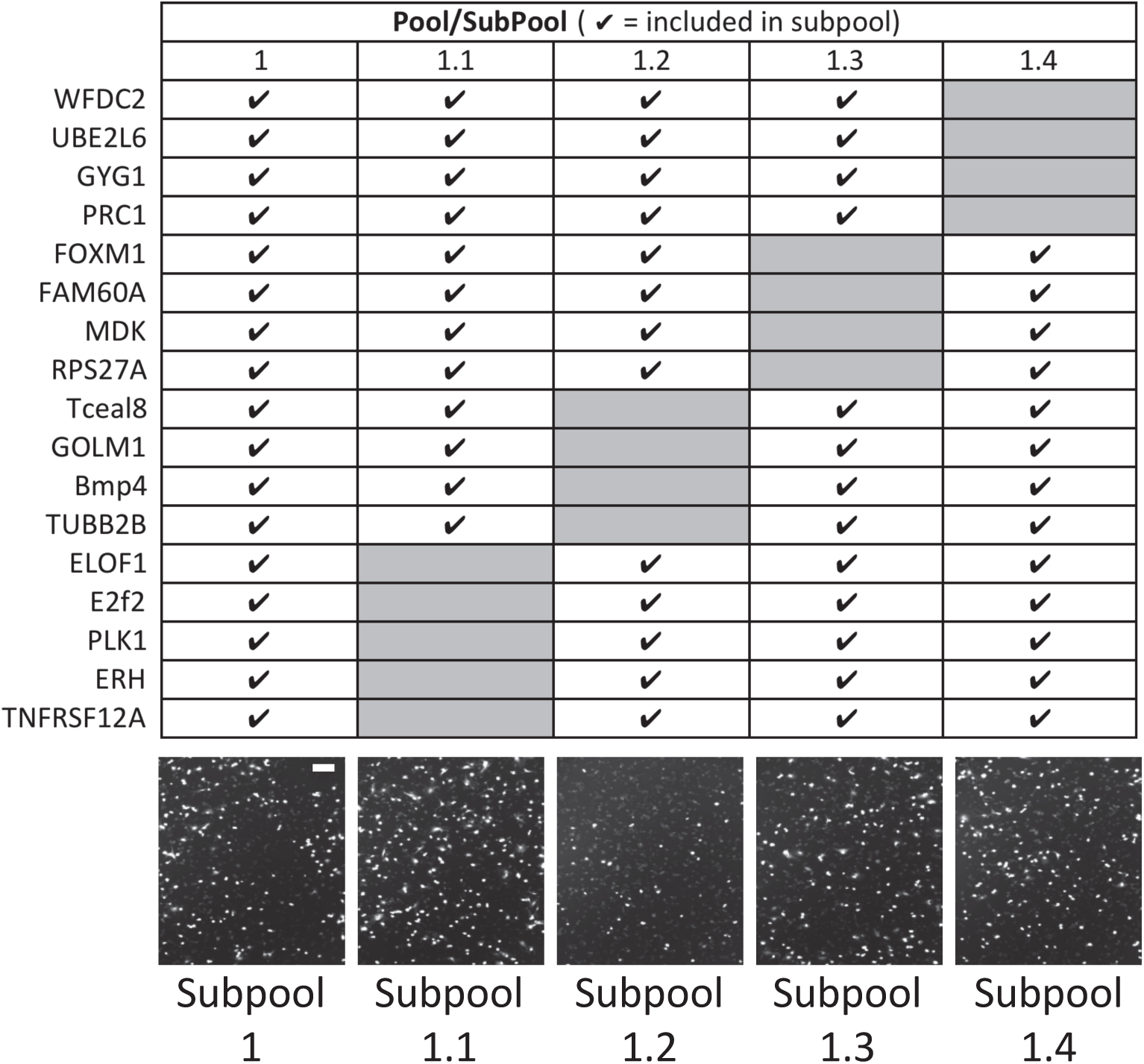
Identification of candidate genes that rescue viability from E2F2-mediated cell death. Each pool contains 12-17 adenoviruses overexpressing indicated genes (indicated with check symbols). Genes removed from each SubPool (1.1, 1.2, etc.) of Pool 1 are indicated by grey boxes. Only gene subpool1.2 was insufficient to rescue viability in the presence of E2F2 overexpression. Scale bars, 200 μm.

**Figure S5.**
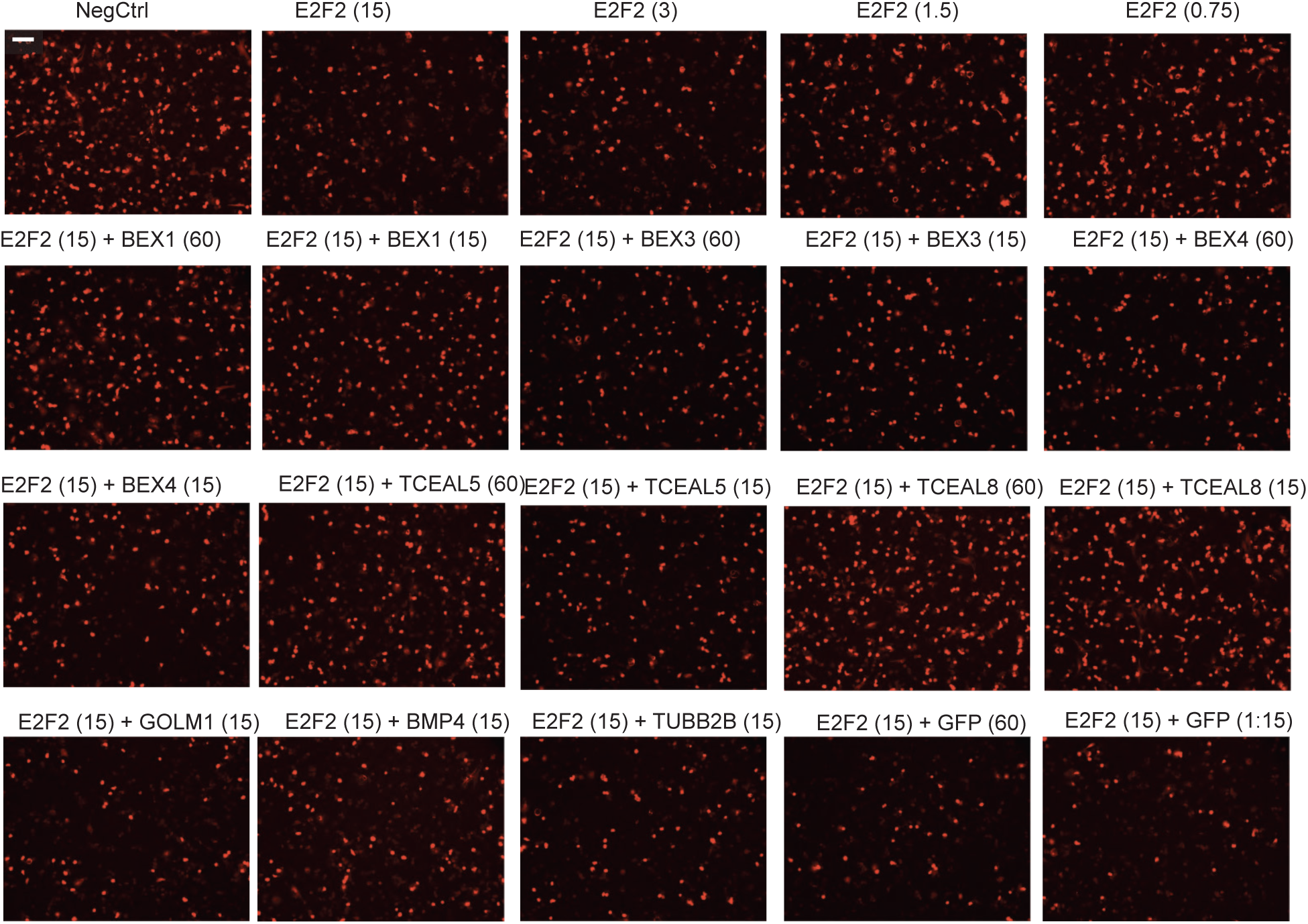
Screening BEX superfamily genes for inhibition of E2F2-mediated cell death. (Top row) E2F2 was expressed by adenovirus vector at various MOI. Cell viability is visualized by the transgenic tdTomato expression (red) at day 7 post-transduction. BEX superfamily genes were co-expressed (MOI 15 or 60) with E2F2 (MOI 15). Note only TCEAL8 and BEX1 exhibit strong inhibition of cell death. MOI is indicated in parentheses. Scale bars, 200 μm.

**Figure S6.**
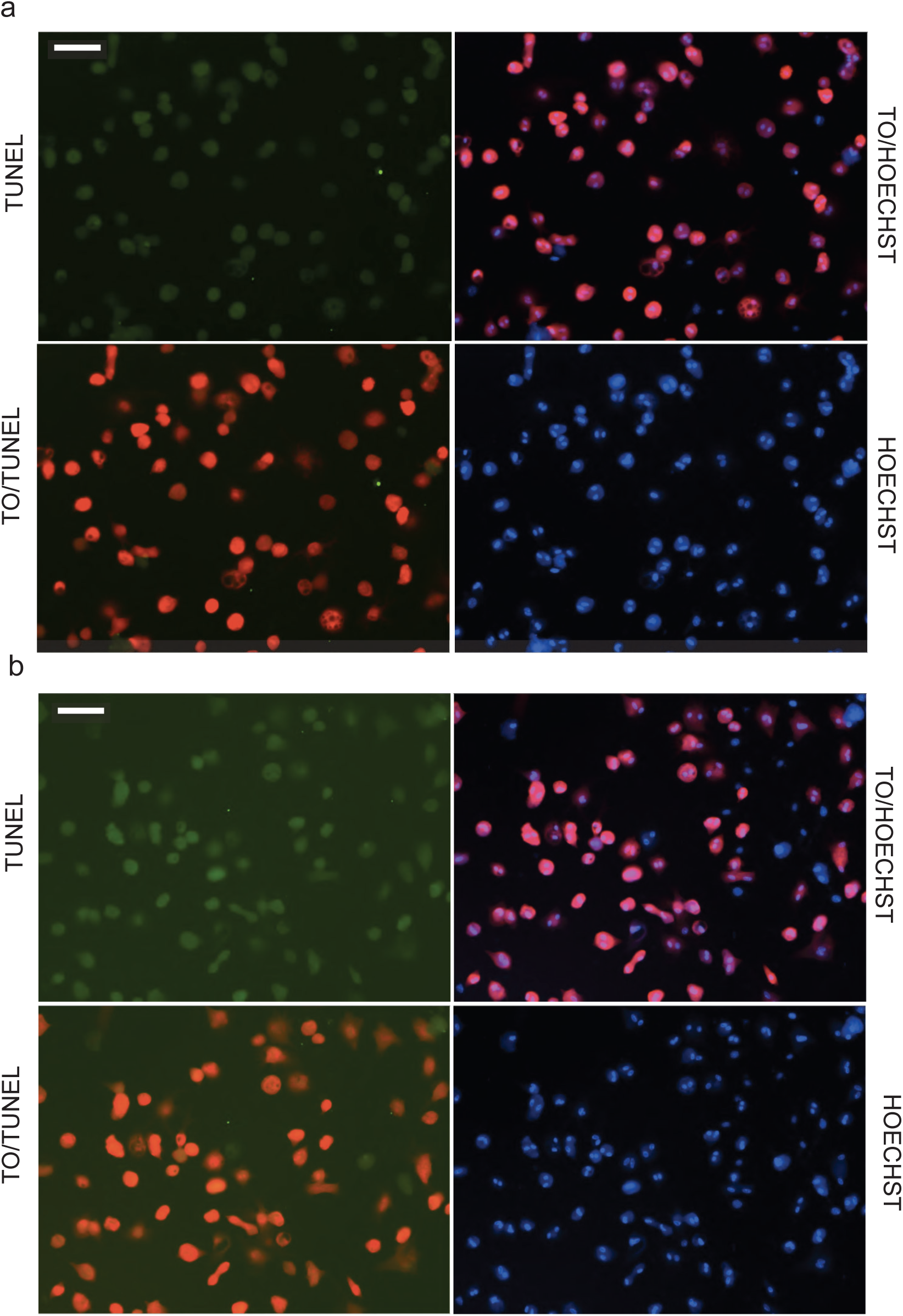
TUNEL staining of E2F2-treated cardiomyocytes co-expressed with E2F2 and TCEAL8 or BEX1. TUNEL labeling shows little genomic degradation at day 5 post-transduction in E2F2-treated cardiomyocytes when co-expressed with TCEAL8 (a) or BEX1 (b). Scale bars, 100 μm.

**Figure S7.**
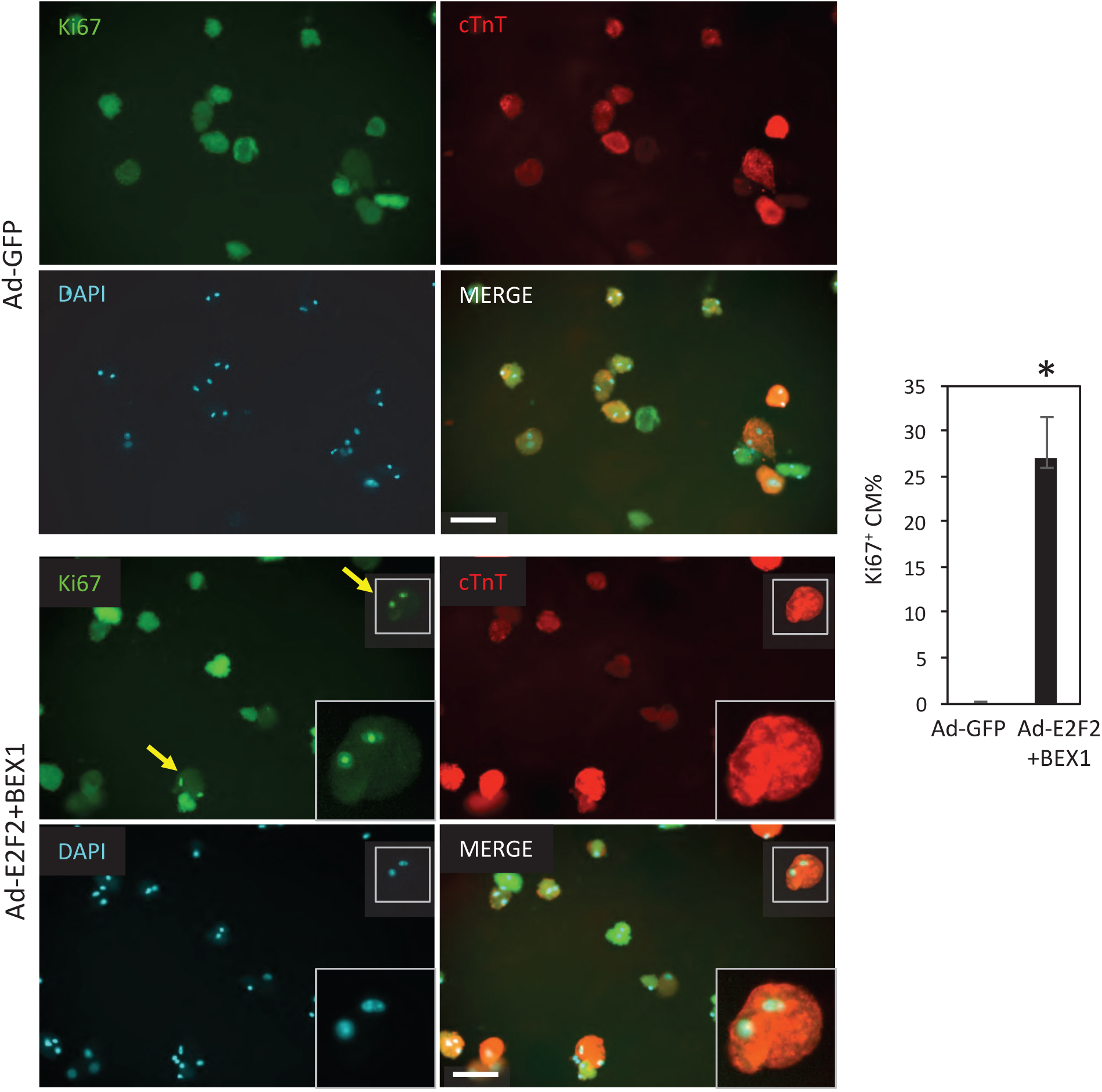
Increase of cardiomyocyte proliferation by adenoviruses E2F2 and BEX1. Adult mouse cardiomyocytes are transduced with viruses at day 1 post-isolation and stained at day 4. Yellow arrows point at Ki67-positive cardiomyocytes. High-magnification views of the boxed areas are shown in the insets. *, p<0.05. Error bars represent standard deviation of mean (n=3). Scale bars, 100 μm.

**Figure S8.**
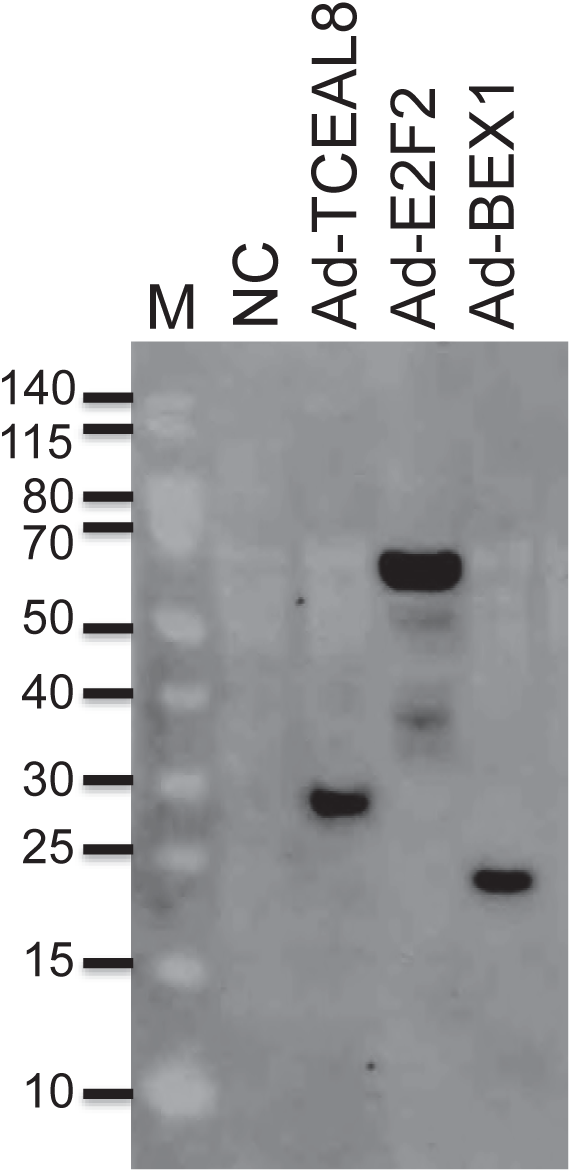
Western blot analysis of V5-tagged E2F2, BEX1, and TCEAL8 protein expression in HEK 293T cells. (a) Expression of E2F2-V5, TCEAL8-V5, and BEX1-V5 proteins were confirmed by western blot analysis of HEK 293T cell lysate at 1 day post-infection. Blot was stained with an anti-V5 antibody. Scale bars, 100 μm.

**Figure S9.**
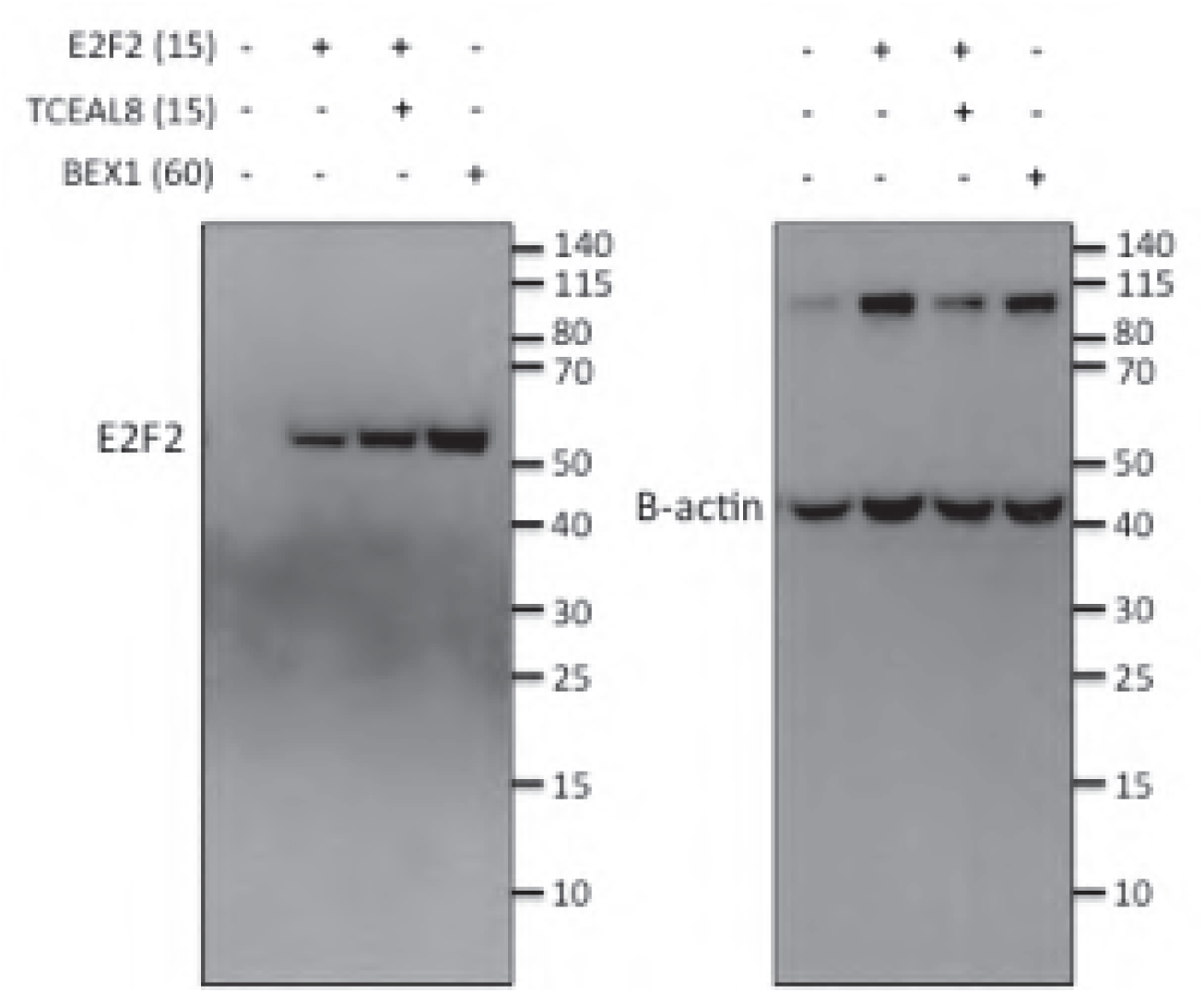
Western blot analysis showing E2F2 co-expressed with Tceal8 and Bex1. Full blot images corresponding to Figure 4c.

**Video S1. Coordinated beating of long-term cultured adult mouse cardiomyocytes.** Cardiomyocytes can be seen beating coordinately at d16 post-isolation. This field of view correlates with the image seen in Figure 1.

